# AER-270 and TGN-020 are not aquaporin-4 water channel blockers

**DOI:** 10.1101/2024.12.04.625365

**Authors:** Lucas Unger, Kim Wagner, Jonas H Steffen, Malene Lykke Wind, Tamim Al-Jubair, Hongjun Zou, Charlotte Clarke-Bland, Rebecca Murray, Bareerah Qureshi, Susanna Lundström, Massimiliano Gaetani, David Poyner, Hoor Ayub, Mark Wheatley, Pontus Gourdon, Andrea J Yool, Susanna Törnroth-Horsefield, Roslyn M Bill, Mootaz M. Salman, Philip Kitchen

## Abstract

Aquaporin-4 (AQP4) is the most abundant water channel protein in the brain. It controls water homeostasis, facilitates glymphatic function and is a drug target for brain edema following injury or stroke. Dysregulation of brain water homeostasis affects millions of people every year leading to death, disability and cognitive decline, for which no medicines are available. Two compounds, AER-270 and TGN-020, are sold as AQP4 inhibitors and a prodrug of AER-270 is currently in a phase I human trial. However, the direct effect of these compounds on AQP4 function has not been unequivocally demonstrated. Our data across multiple cellular and molecular assay systems demonstrate, unexpectedly, that AER-270 and TGN-020 do not inhibit AQP4. Although we observed an apparent inhibitory effect of AER-270 and TGN-020 in the *Xenopus laevis* oocyte assay, there was no effect in assays using reconstituted recombinant AQP4 or mammalian cells expressing exogenous or endogenous AQP4. We identify alternative mechanisms of action for both molecules that may explain previously reported *in vivo* results that were interpreted in the context of AQP4 inhibition. Overall, we conclude that AER-270 and TGN-020 should not be used to investigate the AQP4-dependence of biological processes in the brain.

## Introduction

The human body consists of 60% water. Strict regulation of its water content is required across volume-scales, from the tiny amounts in mitochondrial volume adaption to the multiple liters per day in renal blood filtration. The family of aquaporins (AQPs) are transmembrane proteins found in all kingdoms of life that facilitate passive bidirectional water transport across biological membranes in response to osmotic or hydrostatic pressure gradients. A subset of AQPs additionally transports small polar solutes, including urea, glycerol, hydrogen peroxide, and a small number of AQP proteins also conduct ions (Login and Nejsum, 2023). There are 13 human AQP proteins expressed throughout the body. Aquaporin-4 (AQP4) regulates central nervous system (CNS) water homeostasis and has been identified as a key contributor to dysregulated water homeostasis after traumatic brain injury (TBI), spinal cord injury (SCI) and stroke, which all lead to cytotoxic CNS edema (Manley et al., 2000, Liu et al., 2021).

Each year, millions of people suffer from CNS edema (Jha et al., 2019). It has been estimated that 12 million people per year worldwide suffer a stroke and about 75 million experience TBI or SCI (Dewan et al., 2018, Feigin et al., 2021), with approximately 30% and 60% of these, respectively, developing CNS edema (Jha et al., 2019, Wu et al., 2018). Current treatment options are limited, relying on symptom management, such as osmotherapy (which has limited efficacy), and invasive neurosurgery to reduce intracranial pressure once brain edema has developed. The established role of AQP4 in edema development at the molecular level means it is a validated drug target (Tradtrantip et al., 2017).

AQP4 has also been identified as an important component of the glymphatic system (Iliff et al., 2012), which consists of fluid flow through a CNS-wide network of perivascular spaces to facilitate waste clearance from the CNS parenchyma (Hablitz and Nedergaard, 2021). Exactly how a water channel in the astrocyte endfoot could contribute to the paracellular fluid transport of the glymphatic system is still an open question, but data on *AQP4^-/-^* mice from several laboratories consistently show that AQP4 knockout reduces glymphatic function (Mestre et al., 2018). An AQP4-specific inhibitor would therefore be of great value and allow researchers to begin to investigate how astrocyte endfoot-localized AQP4 could contribute to paracellular glymphatic flow.

Over the last decades, various small molecules were proposed as AQP4 blockers, including antiepileptic agents, such as phenytoin, lamotrigine, and topiramate (Huber et al., 2009a), arylsulfonamides (Huber et al., 2007), loop diuretics such as bumetanide and furosemide (Migliati et al., 2009), and a selection of other chemicals (Huber et al., 2009b). Most of these candidates have been identified using *in silico* methods and showed inhibition of AQP4 expressed in *Xenopus laevis* oocytes with IC_50_ values in the high micromolar range. However, upon testing in mammalian cell culture these compounds failed to show an effect (Yang et al., 2008, Tradtrantip et al., 2017).

TGN-020 and AER-270 (Figure 1a) are commercially available as AQP4 inhibitors. TGN-020 (N-1,3,4-thiadiazol-2-yl-3-pyridinecarboxamide) was initially discovered following a virtual screening approach based on structural similarities to previously reported carbonic anhydrase inhibitors and anti-epileptics (Huber et al., 2009b, Huber et al., 2009a), although such compounds were found ineffective upon re-testing in mammalian cell culture assays (Tradtrantip et al., 2017, Esteva-Font et al., 2016). TGN-020 inhibited osmotic water flux in *Xenopus laevis* oocytes expressing human AQP4-M23 with an IC_50_ of 3.1 µM. A similar study conducted in a different laboratory reported selectivity for AQP4 over other human AQPs (Toft-Bertelsen et al., 2021). Pretreatment with TGN-020 in a middle cerebral artery occlusion (MCAO) mouse stroke model reduced ischemia-induced brain swelling 2-fold (Igarashi et al., 2011) and also improved motor scores and glial scarring in rats 7 days after the inflicted brain damage when administrated post-injury (Pirici et al., 2017). Discrepancies between experiments with TGN-020 and those using *Aqp4*^-/-^ animals highlight a further need for a more comprehensive validation of the inhibitor. For example, TGN-020 had no effect on stimulus-induced extracellular space (ECS) volume dynamics in the *ex vivo* CA1 stratum radiatum (Toft-Bertelsen et al., 2021), whereas the volume change was augmented in *Aqp4*^-/-^ knockout mice (Haj-Yasein et al., 2012). AQP4 knockout was also reported to increase perihematomal edema following intracerebral haemorrhage in mice by approximately 10-fold, whereas TGN-020 had only a minor effect on edema volume (Jeon et al., 2021).

**Figure 1.**
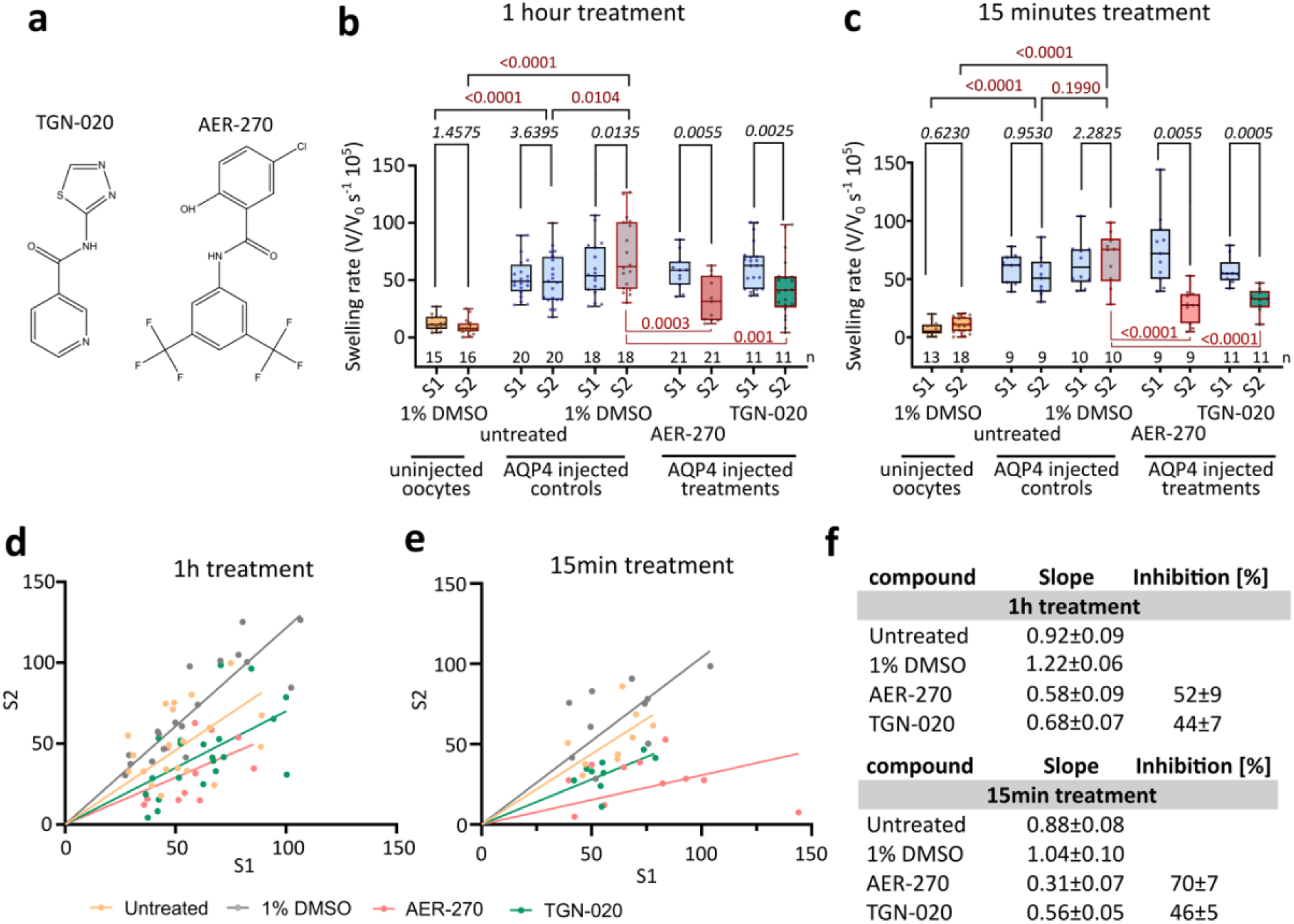
AER-270 and TGN-020 inhibit osmotic swelling of rAQP4-expressing oocytes. (a) Structures of AER-270 and TGN-020 (created using ChemDraw). (b, c) Oocyte swelling rates were measured by time-lapse microscopy upon exposure to a hypoosmotic solution. Data are shown as box plots of three independently performed experiments (total number of oocytes used, n, is indicated above the x-axis). Externally applied 100 µM AER-270 or TGN-020 (both in DMSO) significantly reduced the swelling rate of rAQP4-M1-expressing oocytes after incubation for (b) 1 hour or (c) 15 minutes. (d, e) The swelling rate of each oocyte before (S1) and after (S2) compound treatment following treatment for (d) 1 h or (e) 15 minutes. (f) Calculated slopes and percentage water transport inhibition. Bonferroni-corrected paired t-tests were used to compare swelling rates before and after compound incubation (S1 vs. S2, red p-values). One-way ANOVA with Bonferroni-corrected post hoc t-tests was used to compare effect of compounds to vehicle (black p-values).

AER-270 (N-(3,5-bis(trifluoromethyl)phenyl)-5-chloro-2-hydroxybenzamide) was discovered by screening 500 compounds, leading to identification of the class of phenylbenzamides as having potential AQP4 inhibiting properties using a CHO cell endpoint bursting assay (Farr et al., 2019). IC_50_ values of 0.42 µM, 0.21 µM and 0.39 µM were reported for human, rat and mouse AQP4, respectively, but maximal inhibition varied significantly between species from only approximately 20% (mouse, human) to approximately 70% (rat). Inhibition of AQP1 and AQP5 was also reported. In a water intoxication model, AER-270-treated mice showed a 3.3-fold improved survival rate and injection of the pro-drug AER-271 reduced brain swelling after MCAO by 33% in mice and 62% in rats. A second study reported the prevention of increased brain water content after cardiac arrest in rats (Wallisch et al., 2019). The compound AER-271 was the subject of a phase I clinical trial in the USA to investigate safety and tolerability in 80 healthy volunteers (NCT03804476) but since the trial completion in 2019 no data have been reported. An almost identical trial started in China in 2021 (NCT05200728), which similarly completed in 2023 but has yet to report any data. AER-270 is chemically identical to IMD-0354, a compound used for several decades as a research tool to interfere with NF-κΒ signaling by inhibition of IKKβ (Tanaka et al., 2005, Onai et al., 2004), blockage of which has been demonstrated to reduce CNS water content after MCAO in rats (Li et al., 2016). This highlights the importance of considering off-target effects for these molecules.

Here we present a comprehensive analysis of the inhibitory effects of TGN-020 and AER-270 in *Xenopus laevis* oocytes, mammalian cells overexpressing AQP4, primary human and rat astrocytes, and recombinant human AQP4 reconstituted into proteoliposomes. We investigate direct binding of both molecules to purified AQP4 protein solubilized in both detergent and styrene maleic acid and use several approaches to identify potential off-target effects.

Overall, our data suggest that whilst both TGN-020 and AER-270 appear to inhibit AQP4 in the *Xenopus laevis* oocyte assay, they have no effect on the plasma membrane water permeability of MDCK or HeLa cells overexpressing AQP4, primary human or rat astrocytes, or proteoliposomes containing recombinant human AQP4. Furthermore, we use both hypothesis-based approaches and unbiased target identification to identify several potential off-target effects of both compounds which may explain their *in vivo* effects.

## Results

### AER-270 and TGN-020 inhibit AQP4 when assayed using Xenopus laevis oocytes

The *Xenopus laevis* oocyte assay has been used as the gold standard for determining aquaporin water permeabilities for over 30 years (Preston et al., 1992, Agre et al., 1993). Swelling of oocytes in hypotonic saline solution was therefore recorded using microscopic time-lapse imaging and changes in their cross-sectional area were quantified using ImageJ (representative swelling traces are shown in Figure S1a). The swelling of each oocyte was measured without compound treatment (S1), and after recovery in isotonic saline, the assay was repeated in the presence of the compound or vehicle control (S2), providing a matched pre-inhibition control for each oocyte. An S2/S1 ratio <1 indicates inhibition of water transport (Figure 1). Oocytes expressing rat AQP4 exhibited functional water channel expression as demonstrated by a 4-fold higher swelling rate compared to non-AQP4-expressing (uninjected) control oocytes (Figure 1b). Incubation with 100 µM TGN-020 or AER-270 for 1 hour significantly reduced swelling rates, with 44±7 and 52±9% inhibition, respectively (Figure 1b,d,f). A shorter incubation of 15 minutes had no effect on inhibition by TGN-020, but interestingly AER-270 was more effective at 15 minutes (70±7% inhibition) than at 1 hour (Figure 1c,e,f), possibly suggesting poor stability of the compound in the assay buffer, rapid metabolism of the drug by *Xenopus laevis* oocytes, desensitization of the target of action, or progressive loss of the compound due to partitioning into in an intracellular compartment such as yolk.

### AER-270 and TGN-020 do not inhibit AQP4 in mammalian cells

Although the *Xenopus laevis* oocyte assay is regarded as the gold standard for determining AQP water permeabilities, it is widely acknowledged to suffer from limitations such as inter-laboratory variability and a lack of consistency with other assays of AQP function (Salman et al., 2022, Tradtrantip et al., 2017, Choi et al., 2021). Notably, the lipid composition of *Xenopus laevis* oocyte membranes is significantly different from that of most mammalian cells (Hill et al., 2005). To study the inhibitory effect of AER-270 and TGN-020 in mammalian cells, calcein fluorescence quenching was used (Kitchen et al., 2020) to measure both cell shrinking and swelling in response to hypertonic and hypotonic stimuli, respectively. MDCK and HeLa cells were stably transfected with pDEST47-hAQP4-GFP (Kitchen and Conner, 2015), which increased membrane water permeability by 6-fold and 2-fold, respectively, compared to the parental control lines (Figure 2a,b,g,h; representative raw data shown in Figure S1e–h). Neither AER-270 nor TGN-020 altered cell shrinkage rates at concentrations up to 100 or 300 µM, respectively. AER-270 appeared to decrease the cell swelling rate of AQP4-MDCK cells at 50 and 100 µM (Figure 2b), but we observed significant cytotoxicity at these concentrations (Figure 2e), making interpretation of this result challenging.

**Figure 2.**
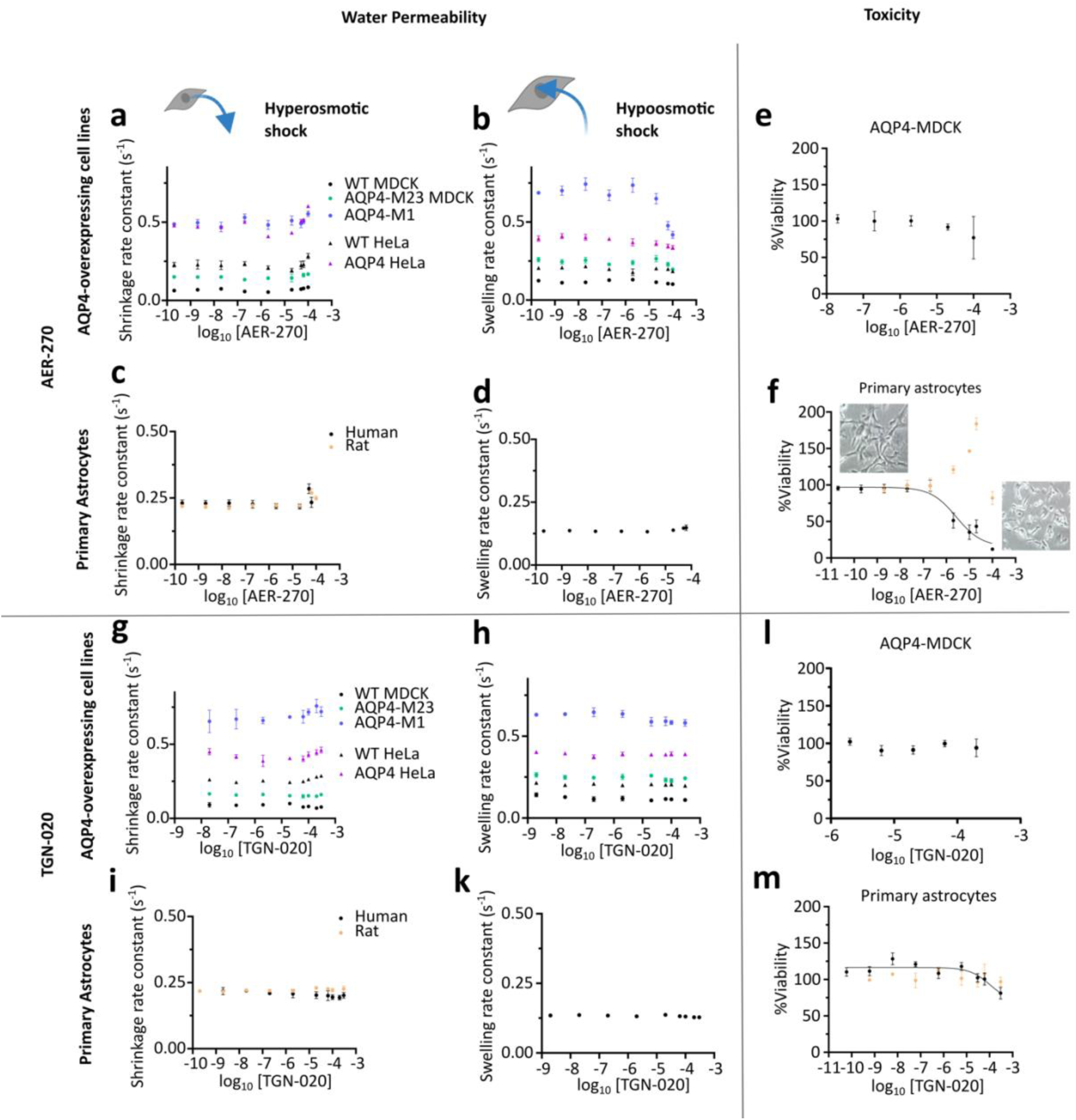
AER-270 and TGN-020 do not affect osmotic swelling or shrinkage rates of mammalian cells. Volume changes of mammalian cells were measured indirectly using calcein-AM following application of a 200 mOsm hyperosmotic gradient (cell shrinking) or 150 mOsm hypoosmotic gradient (cell swelling). Shrinking or swelling rates were measured after 15 minutes pre-treatment with AER-270 (a, b) in MDCK and HeLa cells stably overexpressing AQP4 (n=3-4) and (c, d) primary human and rat astrocytes (n=3-4). AER-270 was cytotoxic in (e) MDCK cells after 30 minutes (n=3) and (f) primary astrocytes after 24 hours (human n=3, rat n=4) measured by MTT assay. (g-k) Cell shrinking/swelling (n=3-4) after 15 minutes TGN-020 treatment and (l,m) cytotoxicity after 6 hours in MDCK cells or 24 hours in astrocytes. Data presented as mean ± SEM.

The full-length AQP4 transcript has two translation initiation sites, leading to two protein isoforms, the 323 residue M1 isoform and the 301 residue M23 isoform. The relative abundance of these isoforms varies between cell types, likely due to differences in expression or activity of RNA-binding proteins (Pisani et al., 2021). Western blotting indicated only the full-length AQP4-M1 isoform was expressed in MDCK cells (Figure S1b-d) stably transfected with full-length AQP4 cDNA, whereas both M1 and M23 were expressed in HeLa cells. To check for isoform-dependent effects, MDCK cells were stably transfected with pDEST47-hAQP4-M23-GFP and the calcein fluorescence quenching assays were repeated. Neither TGN-020 nor AER-270 inhibited AQP4-M23 in these experiments (Figure 1a,b,g,h). We also found no difference in single channel permeability between M1 and M23 after normalising shrinking rates to cell surface abundance (Figure S1j,k).

As the endogenous AQP4 in astrocytes is the desired pharmacological target for both molecules, we tested the agents on both primary human and rat astrocytes. Neither AER-270 nor TGN-020 inhibited water transport in primary human or rat astrocytes (Figure 2c,d,i,k; representative raw data in Figure S1i). AER-270 was highly toxic to human astrocytes in metabolic viability assays using 3-(4,5-dimethylthiazol-2-yl)-2,5-diphenyltetrazolium bromide (MTT), with an LD_50_ of 5.3 µM (Figure 2f). Cell death was confirmed by microscopic inspection. In rat astrocytes, the MTT signal was increased at intermediate concentrations of AER-270, suggesting an effect on cellular redox state or metabolism leading to more rapid NADH-dependent conversion of tetrazolium to formazan. Toxicity was observed only at 100 µM. In contrast, TGN-020 had no effect on MTT signal from rat or human astrocytes at concentrations up to 300 µM (Figure 2m).

A key difference between the assays performed in mammalian cells and other assay systems is the presence of serum in the culture medium. Many small molecules can bind to albumin and other serum proteins, which would reduce their effective concentration. Therefore, the experiments reported in Figure 2 were repeated in AQP4-M1-overexpressing MDCK cells using serum-free cell culture medium. In the absence of serum, we still found no inhibitory effects of AER-270 or TGN-020 on AQP4 (Figure S1l,m).

### AER-270 and TGN-020 do not inhibit AQP4 reconstituted into proteoliposomes

Having observed contrasting results between *Xenopus laevis* oocytes and mammalian cells, we used a proteoliposome-based assay in which the permeability of purified AQP4 protein reconstituted into liposomes is determined by stopped-flow light scattering (Steffen et al., 2022). We used mercury (II) chloride (HgCl_2_) as a control in these experiments. AQP4 has an intracellular mercury ion binding site (Cys178 in loop D; (Yukutake et al., 2008)). Due to the random insertion of AQP4 into liposomes, half of the AQP4 molecules would be expected to have Cys178 exposed to the outside of the proteoliposomes. Since the proteoliposomes used are impermeable to mercury ions, it would be expected that HgCl_2_ inhibits AQP4 by 50% in this system, which is indeed what we observed (Figure 3a). Treatment of AQP4-containing proteoliposomes with 100 µM TGN-020 or AER-270 had no effect on AQP4-dependent water transport (Figure 3a).

**Figure 3.**
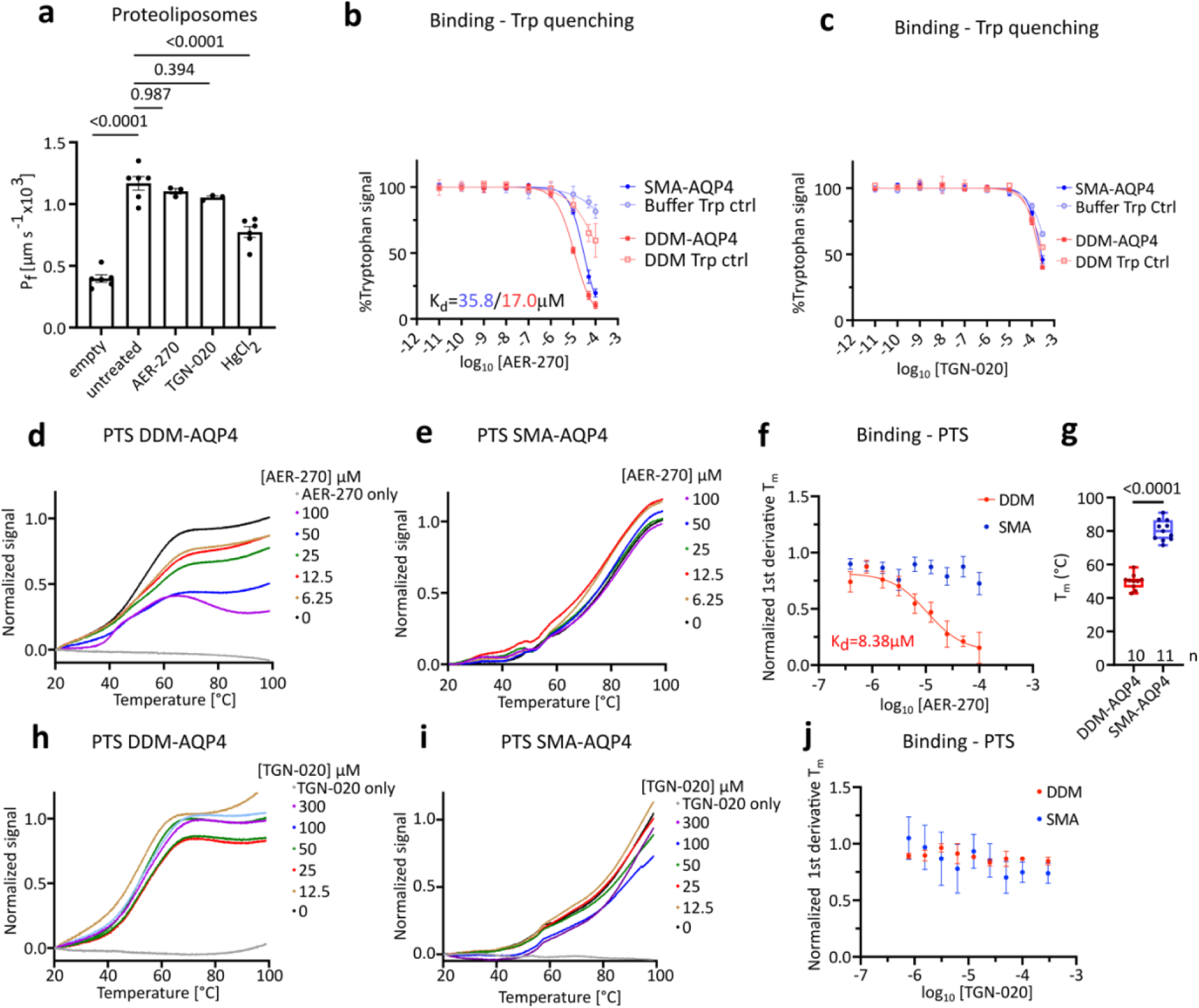
Effects of AER-270 and TGN-020 on recombinant hAQP4 protein. (a) Reconstitution of recombinant hAQP4-M1 into proteoliposomes increased water permeability measured by stopped-flow light scattering. Externally applied 100 µM AER-270 or TGN-020 did not affect hyperosmotic shrinking of AQP4 reconstituted proteoliposomes (n=3). (b, c) Binding to recombinant AQP4 in DDM or SMALPs was evaluated by measuring changes of the protein intrinsic tryptophan fluorescence (Ex 280 nm/Em 336 nm) in the presence of (b) AER-270 (DDM n=3, SMA n=4) or (c) TGN-020 (n=3). Binding affinities (K_d_) were calculated from the compound dose response curve after subtraction of non-specific quenching of free tryptophan (N-acetyl-L-tryptophanamide). (d,e) Representative CPM thermal stability fluorescence traces for (d) DDM-AQP4 and (e) SMA-AQP4 in the presence of AER-270. (f) First derivative of fluorescence intensity with respect to temperature was calculated at the melting temperature (Tm) of the vehicle control (DDM n=5, SMA n=4). (g) SMA-solubilized AQP4 shows an increased thermal stability in comparison to DDM-solubilized AQP4 (DDM n=9, SMA n=7). Melting profiles of (h) DDM-AQP4 (n=3) and (i) SMA-AQP4 (n=4) were measured in the presence of TGN-020 and (j) the change in T_m_ calculated. (a-c, f, g, j). Data are presented as mean ± SEM.

### AER-270 but not TGN-020 binds to recombinant AQP4 with low affinity

To investigate direct binding of AER-270 or TGN-020 to AQP4, we first used tryptophan fluorescence quenching on recombinant hAQP4 produced in *Pichia pastoris* and solubilized using either *n*-dodecyl-β-D-maltoside (DDM) or poly(styrene-*co*-maleic acid) (SMA). To account for fluorescence quenching independent of binding, N-acetyl-L-tryptophanamide was used as a control. AER-270 quenched AQP4 tryptophan fluorescence with an apparent K_d_ of 35.8±4.6 µM for the SMA-solubilised protein and 17.0±3.1 µM for the DDM-solubilised protein (Figure 3b). In contrast, although there was minor quenching of the fluorescence signal at high concentrations of TGN-020, the quenching was identical for AQP4 and N-acetyl-L-tryptophanamide, suggesting a small non-specific effect of TGN-020 on tryptophan fluorescence (Figure 3c).

Notably, AER-270 has intrinsic fluorescence with an emission maximum at 435 nm (Figure S2n) when excited at 280 nm (Figure S2o). The compound fluorescence increased in the presence of the protein (Figure S2p) suggesting either non-radiative energy transfer from a tryptophan residue to AER-270 or a change in the chemical environment of AER-270, both of which would further suggest binding of AER-270 to recombinant AQP4.

Tryptophan fluorescence of a protein will change in the presence of a binding molecule only if the binding results in a change to the chemical environment of a tryptophan residue(s) either by direct close contact or by a conformational change of the protein. Therefore, to supplement the tryptophan fluorescence experiments, we also used a thermal shift assay in which the protein is slowly heated in the presence of a cysteine-reactive fluorescent dye, 7-diethylamino-3-(4’-maleimidylphenyl)-4-methylcoumarin (CPM). As the protein undergoes thermal denaturation, buried cysteine residues become solvent-accessible leading to an increase in the fluorescence intensity of the dye, and small molecule binding is characterized by a shift in the melting profile of the protein, although it has also been suggested that the CPM signal may originate from binding to hydrophobic patches rather than free cysteine residues upon protein unfolding (Wang et al., 2015). Regardless of the origin of the signal, CPM is a sensitive probe of protein unfolding. In this assay, AER-270 altered the melting profile of DDM-solubilized AQP4 in a dose-dependent manner, with an apparent K_d_ of 8.38±0.76 µM (Figure 3d,f), in close agreement with the tryptophan quenching assay. We observed no shift in the protein melting profile for SMA-solubilized AQP4 (Figure 3e,f), possibly due to the enhanced thermal stability of the protein conferred by SMA (melting temperature of 50°C in DDM c.f. 71°C in SMA, Figure 3g). For TGN-020, we observed no shift in the melting profile of either DDM- or SMA-solubilized protein (Figure 3h,i,j). The measured binding affinities for AER-270 are inconsistent with the reported sub-micromolar IC_50_.

### Off-target effects of AER-270 and TGN-020

Rodent edema models have demonstrated positive treatment effects with AER-270 or TGN-020 (Farr et al., 2019, Wallisch et al., 2019, Igarashi et al., 2011, Pirici et al., 2017) but our *in vitro* data suggest that neither compound directly inhibits AQP4. In order to understand these previously reported *in vivo* effects, we investigated potential non-AQP4 targets of both compounds.

Several reported inhibitors of AQP4 in the *Xenopus laevis* oocyte system such as acetazolamide are carbonic anhydrase (CA) inhibitors, suggesting that CA inhibitors might give false positive hits. We therefore tested both TGN-020 and AER-270 in a commercial CA1 assay (Abcam). AER-270 inhibited CA1 with an IC_50_ of 3.3 µM, and an average maximal inhibition of 55±25% (Figure 4a,b). In contrast, TGN-020 did not inhibit CA1 at concentrations up to 300 µM (Figure 4a).

**Figure 4.**
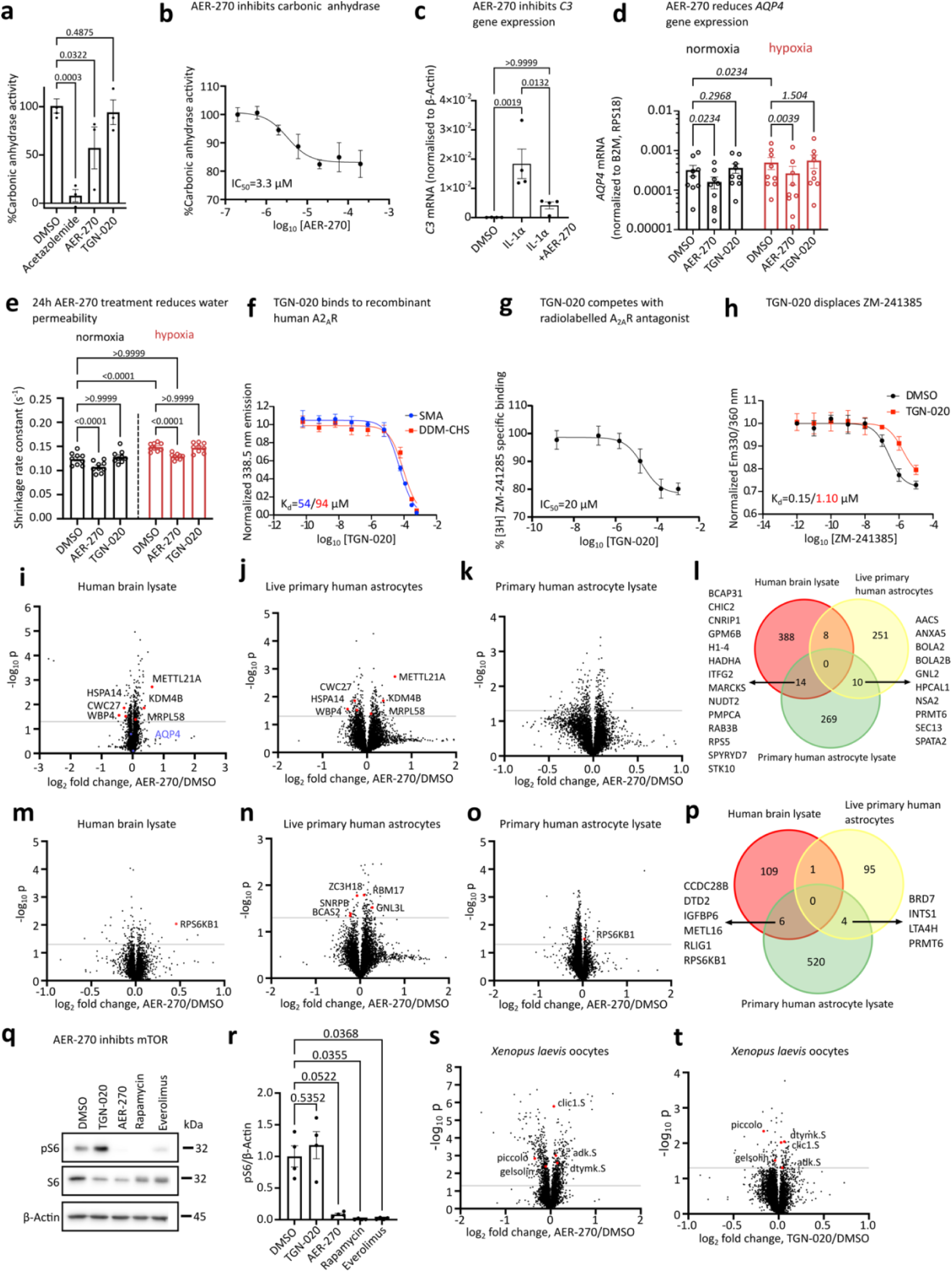
AQP4-independent effects of AER-270 and TGN-020 (a) Inhibition of carbonic anhydrase-1 by TGN-020 or AER-270 was assessed using a commercial enzyme activity kit (n=3). 200 µM acetazolamide was used as a positive control. Data were analysed by one-way ANOVA followed by post hoc t-tests with Holm’s correction. (b) Dose-dependent effect of AER-270 (IC50 = 3 µM). Data shown are normalized to the DMSO vehicle control. (c) Effect of 45 minutes pre-treatment of primary human astrocytes with AER-270 on IL-1⍺-induced upregulation of C3 gene expression (n=4, one-way ANOVA with Bonferroni post hoc test). (d,e) Effect of 24 hours treatment of primary human astrocytes with AER-270 or TGN-020 on (d) AQP4 gene expression (3 donors, two-way ANOVA with Dunnett post hoc test) and (e) osmotic shrinkage rates (4 donors, two-way ANOVA with Bonferroni post hoc test) under normoxic and hypoxic (1% O2) conditions. (f) TGN-020 binds to recombinant human A2AR measured by tryptophan quenching (n=3) and (g) displaces radiolabelled A_2A_R antagonist, ZM-241285 (n=6). A proteome integral solubility alteration (PISA) assay identified proteins that potentially bind TGN-020 or AER-270 in (i, m) human whole brain lysates (5 donors), (j, n) live cultured primary human astrocytes (4 donors) and (k, o) primary human astrocyte lysates (4 donors). (l, p) Hits common to more than one sample type are displayed in Venn diagrams. (q) Representative western blot showing S6 phosphorylation status in primary human astrocytes treated for 24 hours with 100 µM TGN-020 or 200 nM AER-270, or 10 nM rapamycin or everolimus as positive controls and (r) densitometric quantification (4 donors). (s, t) PISA assay identified common targets of AER-270 and TGN-020 in Xenopus laevis oocyte lysates. Data are presented as mean ± SEM.

AER-270 (under the name IMD-0354) is established as an inhibitor of IKKβ; as such it is often used as a tool compound to manipulate NF-κB signalling in cultured cells (Kanduri et al., 2011). However, in the manuscript describing AER-270 as an AQP4 inhibitor, it was reported that AER-270 does not inhibit IKKβ nor any other kinase in three independent kinase panels, although the data were not shown and no details of the assays were provided (Farr et al., 2019). To test the ability of AER-270 to inhibit IKKβ and hence NF-κB signalling in human astrocytes, we induced complement *C3* expression in primary human astrocytes using IL-1, a process known to require NF-κB signalling (Cheng et al., 2017). Stimulation with 3 ng/mL IL-1⍺ for 24 hours increased *C3* gene expression measured by qRT-PCR 198-fold, and pre-incubation with AER-270 at 300 nM for 45 minutes before addition of IL-1⍺ reduced this increase by approximately 85% (Figure 4c), suggesting that AER-270 does inhibit IKKβ, at least in primary human astrocytes.

In order to identify novel targets of AER-270 that might explain its effects *in vivo*, we used SwissTargetPrediction (Daina et al., 2019), an *in silico* tool that predicts protein targets of small molecules based on molecular similarity and a database of known protein-ligand interactions. The top hit for AER-270 was transmembrane serine protease 4 (TMPRSS4). This protein was present in the database due to the inclusion of a 2013 publication describing AER-270 (referred to as compound 2a) as a TMPRSS4 inhibitor with an IC_50_ of 11 µM (Kang et al., 2013). Next, the p38 MAP kinase was predicted as an AER-270 target – interestingly, we and others have previously reported that p38 inhibition in astrocytes reduces AQP4 expression (Salman et al., 2017), suggesting a possible mechanism by which AER-270 could indirectly affect AQP4. To further explore this possibility, we measured *AQP4* transcript expression in primary human astrocytes under normoxia and hypoxia (which is known to increase AQP4 expression (Habib et al., 2014)) following incubation with AER-270 (or TGN-020) for 24 hours. Expression of *LDHA, VEGFA,* and *SLC2A1* (also known as *GLUT1*) were used as positive control hypoxia-responsive genes (Li et al., 2021) to validate the setup (Figure S1q-s). Hypoxia (1% O_2_) for 24 hours increased *AQP4* expression by approximately 2-fold; incubation with 200 nM AER-270 prevented this increase, and also reduced *AQP4* expression under normoxia (Figure 4d). In contrast, TGN-020 at concentrations up to 100 µM had no effect on *AQP4* expression in either condition. The changes in *AQP4* expression with long-term AER-270 treatment reduced water permeability of primary human astrocytes after 24 hours of treatment under both normoxia and hypoxia, correlating with the observed gene expression changes (Figure 4e).

SwissTargetPrediction identified all four adenosine GPCRs (A_1_, A_2A_, A_2B_, A_3_) as potential targets of TGN-020. As our laboratory has extensive experience expressing the human A_2A_ receptor in *Pichia pastoris* (Bawa et al., 2014, Jamshad et al., 2015), we used this recombinant receptor sub-type to study direct interaction with TGN-020 using tryptophan fluorescence quenching. Tryptophan fluorescence of human A_2A_R solubilised in either SMA or DDM supplemented with 1:10 cholesteryl hemisuccinate (DDM-CHS) was dose-dependently quenched by TGN-020 (Figure 4f) with fitted K_d_ values of 54 and 93 µM respectively, in contrast to the minimal quenching previously observed for TGN-020 with AQP4 or N-acetyl-L-tryptophanamide (Figure 3c). We then tested the ability of TGN-020 to displace tritiated A_2A_-selective antagonist ZM-241385 from crude A_2A_-expressing *Pichia* membranes. Competition radio-ligand binding experiments with [^3^H]ZM-241385 (1 nM) as tracer revealed dose-dependent binding of TGN-020 to the A_2A_ receptor (IC_50_ = 20 µM) but with a saturated maximal displacement of only approximately 25% (Figure 4g), suggesting non-competitive binding. We confirmed this for the SMA-solubilised and purified protein using tryptophan fluorescence quenching. Addition of ZM-241385 caused a dose-dependent red-shift of the tryptophan fluorescence spectrum as previously reported (Routledge et al., 2020), which we quantified as the ratio of emission at 330 and 360 nm. This gave a fitted K_d_ of 150 nM for ZM-241385, which was shifted to 1.1 µM by addition of 300 µM TGN-020 (Figure 4h).

We next took a hypothesis-free approach to identifying novel targets of TGN-020 and AER-270 using the protein integral solubility alteration (PISA) assay, a mass spectrometry-based proteomics approach which uses shifts in the aggregation temperature of proteins in the presence of compounds to identify potential targets proteome-wide. We first incubated 100 µM AER-270 or TGN-020 for 10 minutes with primary human astrocyte lysates or human brain lysates to identify direct targets in astrocytes and other brain cell types, respectively. We then separately pre-incubated live human astrocytes for 1 hour with 2 µM AER-270 or 100 µM TGN-020 before lysis to further identify indirect targets whose solubility point changed as a result of activation/inhibition of cell signalling pathways or formation/dissolution of signalling complexes (Gaetani and Zubarev, 2023, Gaetani et al., 2019, Zhang et al., 2022). This approach identified a total of 293 and 530 proteins shifting in solubility upon treatment in astrocyte lysates, and 410 and 116 potential direct targets in brain lysates for AER-270 and TGN-020, respectively (Figure 4i-p). Of these, 14 (AER-270) and 6 (TGN-020) potential targets were conserved between astrocyte and brain lysates, which we considered high-confidence candidate targets of these compounds in astrocytes. In the compound pre-incubation experiments with living cells, we identified 269 and 99 proteins whose solubility was altered by AER-270 or TGN-020, respectively. Pathway analysis of these proteins using gene set enrichment analysis via the Molecular Signatures Database (GSEA MSigDB) was used to identify signalling pathways potentially modulated by AER-270 and TGN-020. Interestingly, the Reactome pathways “*metabolism of RNA*” and “*mRNA splicing*” were among the most significantly enriched pathways for both compounds. STRING analysis identified a spliceosome-associated cluster for AER-270 (CWC27, KDM4B, PPIL1, SF3B3, SMNDC1, SNRNP40, TCERG1, TOE1, WBP4, WDR25), and a different spliceosome-associated cluster for TGN-020 (BCAS2, ERH, GNL3L, RBM17, SNRPB, ZC3H18). In addition, an unfolded protein response cluster was identified for AER-270 (AR, BAG3, HSPA14, HSPA1B, HSPA4, HSPH1, METTL21A, PPID), in agreement with a recent study that reported effects of IMD-0354 on the unfolded protein response in melanoma cells (Feng et al., 2021). We also identified a mitochondrial translation-associated cluster for AER-270 (MRPL58, MRPS11, MRPS12, MRPS34, MTIF3), potentially providing an explanation for the effects of AER-270 that we observed in MTT assays.

The PISA data also identified the ribosomal protein S6 kinase RPS6KB1 as a potential target of TGN-020 in both human astrocyte and whole brain lysates. Furthermore, SwissTargetPredict identified potential binding of RPS6KA1 by TGN-020. As S6 is a key downstream effector of the mTOR signalling pathway, we investigated potential effects of TGN-020 (and AER-270) on mTOR signalling by incubating primary human astrocytes with 100 µM TGN-020 or 200 nM AER-270 for 24 hours and western blotting for phosphorylated S6 (pS6) (Figure 4q). Rapamycin and everolimus both at 10 nM were used as positive controls for mTOR inhibition (MacKeigan and Krueger, 2015). Incubation with TGN-020 caused a minor increase in pS6 to 118% of control (Figure 4r). Surprisingly, AER-270 strongly inhibited S6 phosphorylation by 92%. Previous studies have identified crosstalk between the NF-κB and mTOR pathways via IKKβ phosphorylation of the mTOR inhibitory protein TSC1 (Lee et al., 2007), providing a plausible mechanism-of-action explaining this result.

Finally, we repeated the PISA experiment using *Xenopus laevis* oocyte lysates to identify possible common targets of AER-270 and TGN-020 that might explain the apparent false-positive effect on AQP4-mediated water transport in *Xenopus laevis* oocytes. This identified 968 and 136 proteins shifting in solubility upon exposure to AER-270 or TGN-020, respectively (Figure 4s,t). Of these, 30 were shared between both AER-270 and TGN-020, and included ion channels (clic1), cytoskeleton-modifying proteins (piccolo, gelsolin), and metabolic kinases (adenosine kinase, dTMP kinase). However, due to the relatively poor annotation of the *Xenopus laevis* genome/proteome, several of the identified targets are of unknown function.

## Discussion

Regulation of brain water homeostasis is critical to human health, with its dysregulation implicated in brain edema and impaired glymphatic clearance leading to cognitive decline. While the role of AQP4 in edema formation is well-established, exactly how a water-selective endfoot channel could facilitate bulk paracellular flow in the glymphatic system remains an open question. There is therefore an urgent need for specific and well-validated AQP4 inhibitors in order to understand the molecular mechanisms of glymphatic clearance and interpret existing data.

AER-270 and TGN-020 have beneficial effects in various rodent CNS edema models (Farr et al., 2019, Wallisch et al., 2019, Igarashi et al., 2011, Pirici et al., 2017) and are used to interrogate the AQP4 dependence of molecular and cellular processes in the CNS (Giannetto et al., 2024, Rosu et al., 2020, Harrison et al., 2020). Despite equivocal evidence for their inhibition of AQP4, both compounds are currently advertised and sold as selective AQP4 inhibitors by established chemical vendors (e.g. Sigma-Aldrich, MedchemExpress, and Tocris). While our study reproduced previously reported effects of both compounds in *Xenopus laevis* oocytes, we found no inhibitory effect in mammalian cells expressing exogenous or endogenous AQP4 nor in proteoliposomes containing recombinant human AQP4 protein. Protein binding assays suggested weak binding of AER-270 (but not TGN-020) to AQP4. However, our measured K_d_ values are several orders of magnitude too weak to account for the reported sub-micromolar IC_50_ of AER-270 (Farr et al., 2019).

*Xenopus laevis* oocytes are well-established as an assay system for channel proteins. Numerous exogenous channels have been successfully expressed in the endogenous oocyte background of low channel protein abundance (Lin-Moshier and Marchant, 2013, Zeng et al., 2020). The large physical size of oocyte cells makes them particularly easy to manipulate and observe microscopically. However, significant differences in the efficacies of various drugs against a variety of channel protein targets have been reported in oocytes compared to mammalian cell systems (Cheng et al., 2017, Wang et al., 2014, Dixon et al., 2014, Lacerda, 2001, McIntyre et al., 2001). Additionally, significant differences in membrane lipid composition in the plasma membrane of *Xenopus laevis* oocytes compared to mammalian cells (Hill et al., 2005) may lead to different channel protein conformational ensembles, while differences in protein glycosylation (Hoover et al., 2003, Takacs et al., 2001) and distinct invaginations (Goldin, 2006) in oocytes may also affect drug binding.

Despite a lack of consistency with other assays of AQP function (and despite similar concerns for other channel proteins), the *Xenopus laevis* oocyte assay remains the gold standard for determining AQP water permeabilities (Salman et al., 2022, Tradtrantip et al., 2017, Choi et al., 2021). Here, we identified diverse targets of AER-270 and TGN-020 in *Xenopus laevis* oocyte lysates and we hypothesize that there may be common target proteins or pathways that lead to false positive hits. These include ion channels and cytoskeleton-modifying proteins, both of which could feasibly alter cell swelling rates, although further experiments are required to determine which (if any) of the identified targets could cause false-positive hits.

Off-target effects of putative AQP4 blockers could lead to misinterpretation of *in vivo* data, highlighting a need for the unequivocal validation of direct AQP4 inhibition *in vitro*. To assess the extent of this problem, we performed a literature review of studies using TGN-020 published up to 2024. This identified 58 studies in which TGN-020 was used as an AQP4 inhibitor with no validation that it actually inhibited AQP4-dependent water transport in the system under study. Our data suggest that AER-270 and TGN-020 are not direct AQP4 pore blockers, meaning they likely exert their influence on CNS fluid flow via other target proteins.

Acetazolamide (AZA) has been proposed as an AQP4 inhibitor (Tanimura et al., 2009, Huber et al., 2007). It is established as a carbonic anhydrase inhibitor (Pastorekova et al., 2004), and is used clinically to inhibit carbonic anhydrase to treat a variety of conditions, including glaucoma and idiopathic intracranial hypertension (Pastorekova et al., 2004). In a recent study, rats treated with AZA showed reduced brain water content 7 days after experimental stroke; this was interpreted as being an AQP4-dependent effect (Hao et al., 2022). However, given a recent study suggesting that CSF, not blood, is the source of the initial fluid driving cytotoxic edema (Hussain et al., 2023), it is not surprising that AZA is protective in acute edema models, as it inhibits CSF secretion via inhibition of carbonic anhydrase (Johanson et al., 1992, Dahl et al., 1995). In addition, AZA increases cerebral blood flow (Okazawa et al., 2001), and is used clinically to assess cerebrovascular reactivity (Vagal et al., 2009). This highlights the risk associated with interpretation of data from *in vivo* studies using unspecific and poorly validated “AQP4 inhibitors”. A previous comprehensive study of the inhibitory effect of AZA on AQP4 (similar to what we report here for TGN-020 and AER-270) found no evidence of inhibition (Yang et al., 2008). Interestingly, we also found an inhibitory effect of AER-270 on carbonic anhydrase *in vitro* (Figure 4a).

Understanding of the NF-κB pathway in astrocytes is currently limited. It seems to play a crucial role in astrocyte differentiation (Birck et al., 2021) but astrocytes show limited NF-κB activity under basal conditions (Bhakar et al., 2002, Schmidt-Ullrich et al., 1996). However, neuronal activation, elevated extracellular glutamate (Ghosh et al., 2011) and inflammation (Gupta et al., 2019, Jayakumar et al., 2014) have been reported to activate the NF-κB pathway in astrocytes. Several studies using rodent TBI models have reported that NF-κB activation is associated with cytotoxic cell swelling in astrocytes, which can be inhibited with the anti-inflammatory drug BAY 11-7082 or curcumin (Jayakumar et al., 2014, Li et al., 2016, Laird et al., 2010). IL-1β signaling has been reported to upregulate AQP4 expression via NF-κB in rat astrocytes (Ito et al., 2006). In our experiments, AER-270 was able to block the induction of *C3* expression by IL-1⍺, suggesting that it is indeed able to block NF-κB signaling in human astrocytes, likely via direct inhibition of IKKβ as previously reported. We are not able to account for the report by Farr et al. that AER-270 did not inhibit activity of recombinant IKKβ protein in kinase panels, although it is difficult to assess this result as the data were not shown. Taken together with the finding that AER-270 reduced AQP4 gene expression (Figure 4d), our identification of novel targets for AER-270 suggests that there are a variety of plausible alternative mechanisms-of-action that may explain published *in vivo* findings independent of direct AQP4 pore-blocking.

Having been identified by the virtual screening of molecules with structural similarities to carbonic anhydrase inhibitors and anti-epileptics (Huber et al., 2009b, Huber et al., 2009a), TGN-020 was shown to inhibit water flux in *Xenopus laevis* oocytes expressing human AQP4-M23 with an IC_50_ of 3.1 µM (Huber et al., 2009b). Figure 1b confirms the effect of TGN-020 in oocytes. However, as for AER-270, the oocyte data were inconsistent with the results from mammalian cell lines, primary human astrocytes (Figure 2) and reconstituted, purified human AQP4 (Figure 3). Studies with an MRI PET ligand-labelled TGN-020 reported higher PET signal in wild-type compared to *Aqp4-/-* mice, but labelling was still clearly present in knockouts (Nakamura et al., 2011). It is possible that this binding is to an as-yet-unidentified protein that is differentially expressed in *Aqp4^-/-^* animals. Comprehensive transcriptomic and proteomic analyses of the brains of *Aqp4^-/-^* mice are currently lacking, so it is not possible to compare our list of TGN-020 PISA hits to proteins that are differentially expressed in *Aqp4^-/-^* brains. *Aqp4^-/-^* mice have also been reported to have increased blood-brain barrier permeability (Zhou et al., 2008), but reduced glymphatic function (Mestre et al., 2018). These confounding factors are difficult to control for in PET studies, so differences in the PET labelling intensity may also reflect differences in TGN-020 penetration into and clearance from the brain in *Aqp4^-/-^* mice. Here, we identified adenosine receptors as targets of TGN-020 (Figure 4f,g,h). In rats, caffeine consumption was reported to acutely decrease CSF production, but chronically increase production due to overexpression of the adenosine A_1_ receptor in the choroid plexus epithelium (Han et al., 2009). Adenosine has also been reported to increase cerebral blood flow in rodents (O’Regan, 2005) and humans (Soricelli et al., 1995), and this effect could be reduced by selective antagonism of the A_2A_ receptor. Given that cerebral blood flow and CSF production both contribute to the magnitude of glymphatic flux, our *in vitro* data on the effect of TGN-020 on the A_2A_ receptor and *in silico* prediction of effects on the other three adenosine receptor sub-types suggest a plausible alternative mechanism-of-action by which TGN-020 might exert *in vivo* effects on the glymphatic system.

Based on our data for AER-270 and TGN-020 and the existing literature on a variety of other proposed AQP4 inhibitors, it appears that the *Xenopus laevis* oocyte assay system is susceptible to false positive hits. We strongly recommend that all future studies of putative AQP4 inhibitors supplement *Xenopus laevis* oocyte assays with alternatives such as AQP-reconstituted proteoliposomes and mammalian cell-based assays. We conclude that AER-270 and TGN-020 do not directly inhibit AQP4 water permeability and identify a variety of alternative targets that may explain their effects *in vivo*. The identification of drugs that modulate AQP4 permeability would likely have significant clinical application in the prevention of CNS oedema and enhancement of the glymphatic system – the search for such drugs continues.

## Acknowledgements

L.U. was supported by the European Union Horizon 2020 Research and Innovation programme (Marie Skłodowska-Curie grant agreement No 847419). P.K. is supported by the Biotechnology and Biological Sciences Research Council via a Discovery Fellowship (BB/W00934X/1), and the Aston University RKE Pump Priming Programme. The Aston Institute for Membrane Excellence (AIME) is funded by UKRI’s Research England as part of their Expanding Excellence in England (E3) fund (P.K. and R.M.B.). M.M.S. is supported by a Medical Research Council Career Development Award (MR/W027119/1) and acknowledges support from the BHF Centre of Research Excellence, University of Oxford (grant code: RE/24/130024) and a Biotechnology and Biological Sciences Research Council Pioneer Award (BB/Y512874/1). R.M.B. is supported by a UKRI Frontier Research Grant EP/Y023684/1 (following assessment as an ERC Advanced grant, FORTIFY, ERC-2022-ADG-101096882 under the UK Government Guarantee scheme) and acknowledges a Biotechnology and Biological Sciences Research Council Pioneer Award (BB/Y512874/1). The Chemical Proteomics core facility of the Karolinska Institute (Chemistry I Division, MBB Department), Unit of SciLifeLab and node of the Swedish National Infrastructure for Biological Mass Spectrometry (BioMS), provided full support in the experimental design, performance, and data analysis of the proteomic studies. The authors thank Professor Nanna MacAulay for productive discussions on their experimental approaches.

## Methods

### Materials

TGN-020 was purchased from Sigma-Aldrich and AER-270 from Tocris. [^3^H]ZM241385 ([2-^3^H]-4-(2-[7-amino-2-[2-furyl][1,2,4]triazolo[2,3-a][1,3,5]triazin-5-yl-amino]ethyl)phenol), specific activity 80 Ci/mmol, was purchased from American Radiolabelled Chemicals (Royston, UK).

### Mammalian cell culture

The mammalian cell lines were cultured in T75 flasks (Corning) with Dulbecco’s modified Eagle’s medium (DMEM, Gibco) supplemented with 10% fetal bovine serum (FBS, Sigma-Aldrich) in a humified incubator at 37°C and 5% CO_2_ without antibiotics. Cells were routinely subcultured at ∼90% confluency using 0.25% trypsin-0.53 mM EDTA (Thermo Fisher). For primary rat astrocytes DMEM supplemented with 20% FBS was used for culturing and accutase (Thermo Fisher) served as the cell dissociation reagent. Primary human cortical astrocytes were purchased from Sciencell (cat no. 1800), maintained in astrocyte medium (Sciencell cat. no 1801) and subcultured using accutase at ∼90% confluency for up to 5 passages. Astrocyte identity was confirmed by immunocytochemistry for EAAT1, S100B and GFAP. All cells were routinely tested for mycoplasma using EZ-PCR mycoplasma detection kit (Biological Industries).

### MTT toxicity assay

Cells were seeded into transparent 96well plates and grown to a density of ∼80-90% for at least 24 hours before starting the experiment. Cell media was then replaced by 100 µL fresh media containing the compounds that needed testing for their toxicity. After the indicated incubation time, 100 µL media containing 0.5 mg/mL MTT reagent (thiazolyl blue tetrazolium bromide, Sigma-Aldrich) was added on top and after 1 hour (HEK293T)/2 h (MDCK)/3 h (primary astrocytes) incubation at 37 °C, the cells were washed once with pre-warmed PBS and then 100 µL DMSO (Sigma-Aldrich) was added. After 20-30 minutes light-protected shaking, the absorbance at 570 nm was measured using a plate reader (Fluostar Omega, BMG). The optical density signal of compound-treated cells was normalised to the respective vehicle control. The assay was conducted in technical triplicates.

### Transfection of mammalian cells

Transient transfections were conducted using polyethyleneimine (PEI, branched, Sigma-Aldrich) in a 6:1 ratio with plasmid DNA. Stably transfected aquaporin cell lines were created using pDEST47 expression vector with neomycin resistance and C-terminal GFP-tag (Kitchen et al., 2015). 24 hours after cells were transiently transfected in 6-well plates, G418 was added (700 µg/mL for MDCK, 500 µg/mL for HeLa) until clear resistant colonies were formed. Clones were picked using trypsin in a micropipette tip, transferred into 96 well plates and then to T25 flasks. Clones were screened for AQP expression using calcein quenching and western blotting to select clones with high expression.

### Cell surface biotinylation

Cell surface biotinylation was performed as previously described. Briefly, AQP4-GFP-transfected cells were labelled using EZ-Link Sulfo-NHS-SS-Biotin (Thermo Fisher) at 0.33 mg/mL in PBS, followed by quenching of excess reagent with 50 mM glycine. Cells were lysed in 120 µL tris-triton lysis buffer (1% v/v triton X-100, 100 mM NaCl, 2 mM MgCl_2_, 25 mM, tris pH 7.4) containing protease inhibitor cocktail (cOmplete, Merck) for 45 minutes. Lysates were centrifuged at 18000xg at 4°C for 10 minutes to remove cell debris and the supernatant added to neutravidin-coated 96well plates (Pierce) for 2 hours at 4°C, followed by blocking with 3% BSA in PBS for 1 hour at RT. The plate was incubated with primary antibody (anti-GFP, Abcam, ab6556) overnight at 4°C, followed by HRP-conjugated secondary antibody (CST) at RT for 1 hour and o-phenylenediamine dihydrochloride (OPD, Sigma-Aldrich) until a signal appeared. Absorbance was measured at 450 nm using a Fluostar Omega plate reader (BMG). The assay was performed in technical triplicates.

### Recombinant protein expression and purification

*Pichia pastoris* X33 was used as the expression system, and transformed with pPICZB-AQP4 or pPICZB-A_2A_R. Yeast were grown on YPD plates containing 0.1 mg/mL zeocin (Thermo Fisher) at 30°C for three days and single colonies picked for 50 mL BMGY starter cultures (20 g/L peptone, 10 g/L yeast extract, 100 mM potassium phosphate buffer pH 6.0, yeast nitrogen base (YNB), 0.4 µg/mL biotin, 0.5% v/v glycerol, 0.1 mg/mL zeocin) at 30°C in baffled flasks shaking at 220 rpm. After 24 hours, 2 mL of starter culture was inoculated into a 2 L Applikon bioreactor vessel containing 1L BMGY with proportional-integral-derivative control set to maintain pH at 6.0, temperature at 30°C, and dissolved oxygen at 30% of maximum. Upon starvation (determined by dissolved oxygen spike), a glycerol fed batch was initiated using 50% v/v glycerol for four hours at 14 mL/hour. Once the glycerol was consumed (determined by dissolved oxygen spike), protein expression was induced using a 50% v/v methanol feed added at at 4.8 mL/hour for 36–48 hours at either 30°C (AQP4) or 22°C (A_2A_R). For A_2A_R, DMSO was added to a final concentration of 2% v/v during induction. Cells were harvested in breaking buffer (5% glycerol, 2 mM EDTA, 100 mM NaCl, 50 mM sodium phosphate buffer pH7.4) supplemented with cOmplete protease inhibitor cocktail (Merck) and homogenised using an EmulsiFlex C3 (Avestin). The cell lysate was centrifugated at 4,000xg for 20 minutes to remove unbroken cells and cell debris, followed by ultracentrifugation of the supernatant at 1500,000xg for 90 minutes to obtain membranes. The membranes were resuspended in membrane resuspension buffer (20 mM Tris-HCl pH8, 20 mM NaCl, 10% glycerol) at a concentration of 180 mg/mL and either stored at -80°C or used immediately. For SMA solubilisation membranes were diluted 1:3 with solubilisation buffer (20 mM Tris-HCl pH8, 300 mM NaCl) containing 3.33% SMA (final concentrations SMA 2.5%, membranes 45 mg/mL) and incubated at RT for 1 h. The SMA2000 was produced in-house according to a published protocol (Rothnie, 2016). For DDM solubilisation, 8% DDM in solubilisation buffer was added 1:1 to the membrane and incubated for 2 h at 4°C. The soluble fraction was isolated by ultracentrifugation at 100,000xg for 45 minutes. Solubilised proteins were incubated at 4°C with Ni-NTA resin (1/10 (v/v), Qiagen) supplemented with 10 mM imidazole to minimise non-specific binding, either overnight (SMA) or for 2 hours (DDM). Immobilized metal affinity chromatography (IMAC) was performed using gravity flow columns. Resin was washed with 100X resin volume of 20 mM imidazole in solubilisation buffer. For AQP4 this was followed by 100X resin volume of 75 mM imidazole. Proteins were eluted using 200 mM (A_2A_R) or 300 mM (AQP4) imidazole in solubilisation buffer. For the DDM-solubilised proteins, all solutions were supplemented with 0.03% w/v DDM (with 1:10 cholesteryl hemisuccinate for A_2A_R). Eluted proteins were concentrated using ultrafiltration (Vivaspin concentrator with 50 kDa MWCO, Sartorius), and further purified using size-exclusion chromatography (superdex200 10/300 GL on Äkta Pure system). The final protein purity was assessed by SDS-PAGE and protein concentration was quantified using a BSA standard curve on the same gel.

### Western Blotting

All polyacrylamide gels were hand casted (4% separating gel pH8.8, stacking gel pH6.8 percentage indicated in Figures) and 0.1% (w/v) SDS. Bio-rad protein electrophoresis/blotting equipment was used. The samples were incubated with Laemmli buffer for 10 minutes at room temperature before loading into the gel and running it using Tris/glycine buffer (25 mM Tris pH8.8, 192 mM glycine, 0.1% SDS) at constant voltage of 180 for 60 minutes or until the dye front reached the gel bottom. For further immunoblotting proteins were transferred from gel onto a PVDF membrane in transfer buffer (25 mM Tris pH 8.8, 192 mM glycine, 20% MeOH) at 100 V for 60 minutes. Membranes were blocked with 10% (w/v) skimmed milk powder in 0.1% PBS-T for 1 h at RT. Primary antibodies were incubated with 5% BSA overnight at 4°C, followed by the HRP-conjugated secondary antibody at RT for 1 h, before applying enhanced chemiluminescence reagents (Thermo Fisher). The protein signal was detected using a digital imaging system. For molecular weight analysis PageRuler Plus Prestained Protein Ladder (Thermo Fisher) was used.

For blue native PAGE, whole cells were lysed in native lysis buffer (20 mM bis-tris, 500 mM aminocaproic (aminohexanoic) acid, 20 mM NaCl, 2 mM EDTA, 10% glycerol, 0.5% v/v triton X-100, pH 7). For electrophoretic separation, the cooled tank was filled with blue cathode buffer (50 mM tricine, 15 mM bis-tris, 0.02% w/v Coomassie G250, pH 7) in the inner tank and anode buffer (50 mM bis-tris, pH 7) in the outer reservoir. Samples were loaded into 4–8% hand-cast gradient gels without separate stacking gel and separated in a cooled tank at 100 V for 15 minutes followed by 1 h 45 minutes at 180 V. NativeMark™ Unstained Protein Standard (Thermo Fisher) served as a size marker.

### Stopped-flow spectroscopy with proteoliposomes

To evaluate the activity of AQP4 in the presence of channel-blockers, we characterized water transport using a proteoliposome assay (Steffen et al., 2022). For this purpose, full-length human AQP4 was produced in *Pichia pastoris* and purified in octyl glucoside (OG, Anatrace) as previously described (Al-Jubair et al., 2022). Following purification, AQP4 was reconstituted in liposomes with a lipid composition of 1-palmitoyl-2-oleoyl-sn-glycero-3-phosphocholine (POPC), 2-oleoyl-1-palmitoyl-sn-glycero-3-phospho-rac-(1-glycerol) (POPG) and cholesterol in a 2:1:2 ratio (Tong et al., 2012). Specifically, POPC, POPG and cholesterol (Sigma) were mixed in a 2:1:2 ratio and dissolved in chloroform in a glass vial to a concentration of 25 mg/mL followed by dehydration using N_2_ forming a thin lipid layer. Upon dehydration, the lipid film was kept under a light N_2_ stream for additional 2-4 hours to achieve complete chloroform removal. The lipid film was rehydrated with reconstitution buffer (20mM HEPES-NaOH pH 8.0, 200 mM NaCl) with 10 mM (5)6-carboxyfluorescein (Sigma), to a concentration of 20 mg/mL lipids. The lipid suspension was sonicated in a sonication bath for 3×15 minutes, with a 5 minute break between cycles. The lipids were frozen in liquid nitrogen and thawed three times. When thawed for the third time, the lipids were passed through a 100 nm polycarbonate filter 11 times, using an extruder (Mini-Extruder, Avanti). The lipids were diluted to 4 mg/mL with reconstitution buffer containing 25% glycerol and 1% OG, after which 10% Triton X-100 was added to the sample to a final concentration of 0.02%. AQP4 was added to the lipid suspension using a lipid-to-protein-ratio (LPR) of 200 and each sample was dialyzed overnight at 4°C in reconstitution buffer. The samples were centrifuged at 57,000xg for 1.5 hours and the resulting pellets were re-suspended in reconstitution buffer. Inhibitors (100 µM TGN-020 and AER-270, 1mM HgCl_2_) were added during resuspension in reconstitution buffer and incubated for 1 h on ice before measuring activity. The shrinkage assay was performed on an SX-20 Stopped-Flow Spectrometer system (Applied Photophysics), where the liposomes were mixed with reaction buffer with 200 mOsm sucrose.

Data were collected at 495 nm at a 90° angle for 2 s. All data were collected at 18°C. Empty liposomes were used as a negative control. Data for each sample were the average of 10 readings. Data were fitted using a double exponential fit. The smallest rate constant is unaffected by changes in AQP4 reconstitution efficiency, while the larger rate constant corresponds liposomes containing AQP4. This rate constant (k) was used to calculate the osmotic water permeability, P_f_ (cm/s) = k / ((S/V_0_)*V_w_*(C_out_-C_in_)), where (S/V_0_) is the initial surface area to volume ratio of the liposome, V_w_ is the partial molar volume of water (18 cm^3^ mol^-1^), C_in_ and C_out_ are the initial concentrations of solute on the inside and on the outside of the vesicles, respectively. The assay was conducted in technical triplicates.

### Measurement of osmotic AQP permeability in *Xenopus laevis* oocytes

Unfertilized oocytes were harvested from female *Xenopus laevis* and prepared as described previously (Goldin, 1992) in accord with approved protocols for animal care and handling (University of Adelaide Animal Ethics Committee; #M-2022-050 AEC). Oocytes were injected with 50 nL of water containing 1 ng of rat AQP4 (Gene ID25293) cRNA and incubated for two or more days at 18°C in Frog Ringers saline (96 mM NaCl, 2 mM KCl, 5 mM MgCl_2_, 0.6 mM CaCl_2_, 5 mM HEPES, pH 7.6) containing 5% (V/V) horse serum, 100 U/mL penicillin, 0.1 mg/mL streptomycin and 5 mg/mL tetracycline as described previously (Chow et al., 2021). For the experiments, isotonic sodium saline (96 mM NaCl, 2 mM KCl, 5 mM MgCl_2_, 5 mM HEPES, pH 7.6, without added Ca^2+^), and 50% hypotonic saline (isotonic saline diluted with an equal volume of Milli-Q purified water) were used.

Osmotic swelling was performed at room temperature under a dissecting microscope mounted with a Cohu CCD video camera; time-lapse images were captured using NIH ImageJ software (https://imagej.nih.gov/ij/). For osmotic swelling measurements, oocytes were transferred from isotonic saline to 50% hypotonic saline, and images were captured at 1 Hz for 30 seconds. After the first swelling assay (S1), the oocyte was removed immediately, rinsed in isotonic saline, and then incubated for 2 h in isotonic saline containing: no added compound (saline control); 0.1% DMSO (vehicle control); or candidate AQP modulators (100 μM TGN-020, 100 μM AER-270). After incubation, the second osmotic swelling measurement (S2) in 50% hypotonic saline was carried out in the continued presence of the same treatment. Double swelling assays allowed each oocyte to serve as its own control, minimising effects of differences in expression levels between oocytes.

Swelling rates were quantified from plots of oocyte volume (standardized to the initial volume) as a function of time after introduction into the hypotonic saline at time zero, fit by simple linear regression to measure slope values. To assess effects of treatments across all oocytes, the initial swelling rates (S1) for each oocyte were plotted against the post-treatment swelling rate (S2). Fit by linear regression, the slopes of the S1-vs-S2 analysis showed the overall effect of the treatment, without influence of differences in channel expression between oocytes. For S1 vs S2 plots, a slope value near 1 indicates no difference between the initial and treatment conditions; a slope value < 1 indicates an inhibitory effect of the treatment; and a slope value > 1 indicates potentiating effects of the treatment on water flux.

### Calcein fluorescence quenching

24 h before the experiment cells (6×10^3^ MDCK, 8×10^3^ HeLa per well) were seeded into a black 96-well clear-bottom plate. Primary astrocytes were seeded from an 80% confluent flask and the experiment conducted when the cells reached similar confluency again. The cells were loaded with 5 µM of calcein-AM (Thermo Fisher) for 90 minutes at 37°C and after washing, covered with 75 µL HEPES-buffered DMEM containing different inhibitors. All media were supplemented with 0.75 mM probenecid to prevent calcein export. After 15 minutes incubation with the inhibitors, the plate was transferred to a pre-heated (37°C) plate reader (Fluostar Omega, BMG) and the measurement conducted in well-scan mode. 400 mM mannitol in cell culture medium (shrinkage) or distilled water (swelling) was injected to generate an osmotic gradient and the fluorescence intensity (Ex485, Em525) measured once every 60 ms for 20 s. Water permeability was quantified by calculating the rate constant k through kinetic curve fitting (shrinkage C+A*exp(-k*x), swelling C-A*exp(-k*x)). After the experiment, the cells were observed by phase contrast and fluorescence microscopy to check for changes in cell morphology indicative of toxicity.

### Tryptophan quenching

Intrinsic protein tryptophan fluorescence was measured with a Perkin Elmer LS 55 fluorescence spectrometer in a 3 mm quartz cuvette (high precision cell, Hellma Analytics) using 100 µL of 50 µg/mL protein (corresponding to an AQP4 monomer concentration of 1.5 µM) in 20 mM Tris pH 8.0, 300 mM NaCl. The sample was excited at 280 nm and the emission spectrum scanned from 290–500 nm. Binding affinities were acquired by successively adding 1 µL of compound at 10-fold increasing concentrations and averaging six consecutive scans for each concentration. Data were normalized to corresponding DMSO controls to account for dilution and solvent effects. To check for non-specific quenching, N-Acetyl-L-tryptophanamide (Sigma) was used at an empirically determined concentration that gave the same magnitude of fluorescence intensity as the protein. For intensity quenching, the intensity at the emission maximum was used for analysis. For peak redshift, the ratio of emission at 330 and 360 nm was calculated.

### Protein thermal shift assay

The 5 mg/mL CPM (7-Diethylamino-3-(4’-Maleimidylphenyl)-4-Methylcoumarin), Thermo Fisher) stock was diluted into the protein buffer at a concentration of 180 µg/mL (detergent) or 36 µg/mL (polymer), thoroughly mixed and light-protected incubated for 60 minutes at room temperature. Protein (final 35 µg/mL), tested molecules and 10-fold diluted CPM dye mixed and 50 µL per sample transferred into a real-time PCR 96well plate (Primer Design). The measurement was conducted in a Lightcycler 480 instrument (Roche). After an initial equilibration to 20 °C, the temperature was increased up to 99 °C with a ramp rate of 3.6 °C per minute while monitoring the fluorescence with the SYBR Green filter (Ex465 nm/Em510 nm). The data was analysed by calculating the first derivative of the fluorescence signal against the temperature. The melting point TM for the protein in the absence of the tested molecules was determined at the inflexion point of the fluorescence trace by calculating the maximum of the Lorentizian-curve-fitted first derivative. Changes in thermal stability in the presence of the tested molecules were calculated by plotting the normalised fluorescence signal at the T_m_ of the pure protein. The experiment was conducted in technical quadruplicates.

### Measurement of gene expression

Cells were seeded in a 6well plate and treated once they reached a confluency of ∼80% with one well per condition. After 24 h, the cells were washed with cold PBS, and the RNA was extracted according to the instructions of the Meridian Isolate II RNA Mini Kit (SLS). For the hypoxia-treated cells, all steps up to the lysis were performed in the hypoxic glove box (Coy Laboratory Products). The RNA concentration was measured with a Nanodrop Spectrophotometer and examined for integrity by gel electrophoresis before continuation with reverse transcription (QuantiTect Rev. Transcription Kit, Quiagen). Quantitative PCR (qPCR) was conducted using SYBR Green master mix (PrecisionPLUS SY qPCR, PrimerDesign) on the LightCycler II instrument (Roche).

Relative gene expression was calculated using the Pfaffl method (Pfaffl, 2001). Primer efficiency was evaluated with a cDNA 2-fold dilution series. Primer sequences are listed in Table 1. *B2M* and *RSP18* were used as housekeeping genes.

**Table 1.**
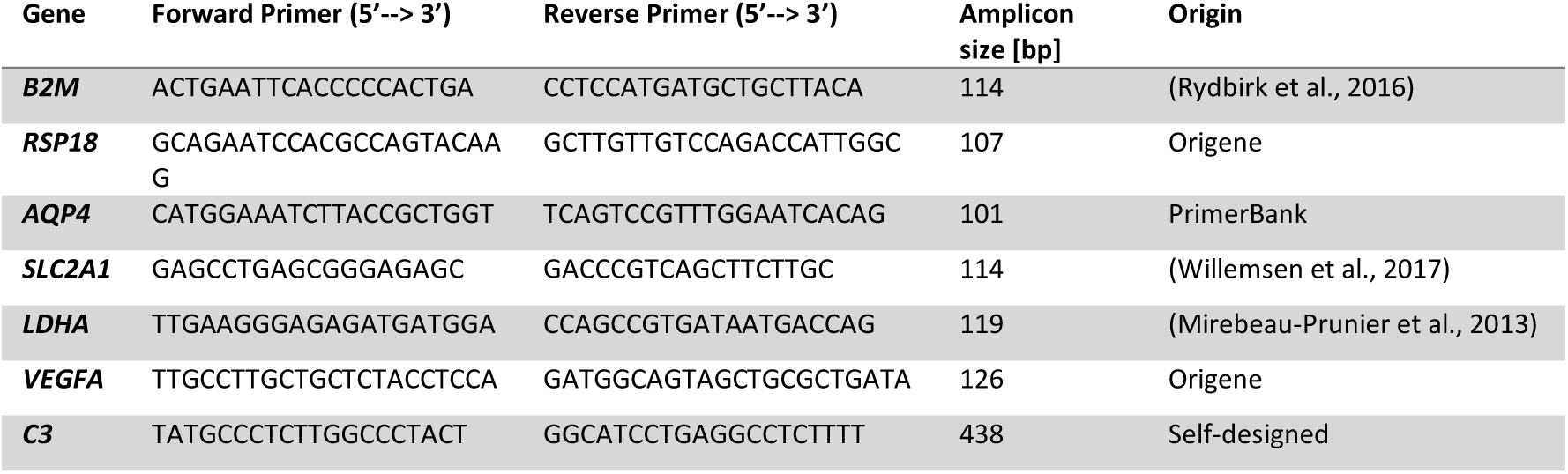
qPCR primer pairs. The table shows the details of the used primer pairs to detect human gene expression levels. Primer sequence, resulting amplicon size, and origin/reference are listed.

### Carbonic anhydrase activity

A colorimetric screening kit (Abcam, ab283387) was used accordingly to the supplier’s protocol. Briefly, assay buffer, recombinant carbonic anhydrase-1 enzyme and the tested compounds were added to wells of a transparent 96-well plate. After adding carbonic anhydrase substrate and mixing, the absorbance at 340 nm was immediately measured once per minute for 60 minutes at 37°C. The background of enzyme-independent substrate conversion was determined using wells with substrate and buffer but no enzyme. The enzyme activity was calculated by linear fitting to the first 20 minutes of the absorbance timeseries, and the enzyme-free background was subtracted. The assay was conducted in technical triplicates.

### Proteome integral solubility alteration (PISA) assay

The protein thermal stability-based assay to identify potential compound binding proteins was performed according to a published protocol (Gaetani and Zubarev, 2023) by Dr Massimiliano Gaetani and his team at the Karolinska Institute in Stockholm, Sweden. The experiment was conducted after compound treatment of either cell lysates (*Xenopus laevis* oocytes, human astrocytes, whole brain lysate) in 0.4% NP40 PBS (Sigma-Aldrich) or living human astrocytes (ScienCell) in technical triplicates. *Xenopus laevis* oocytes were purchased from Xenopusresource and defolliculated according to an established protocol (Anthony et al., 2000) and, after washing, resuspended in the lysis buffer. Primary human astrocytes (ScienCell) of four donors were grown in separate cell culture flasks to 90% confluency and then pooled with equal cell amounts before cell pellet was prepared for the experiment in cell lysate or aliquoted in cryo-vials for the later compound treatment in cultured cells. Whole brain lysate from five different donors (NB820-59177, Novus Biologicals) was pooled and buffer-exchanged before the experiment. All cell lysates were split into 800 µL samples of 1 µg/µL total protein per replicate and treated with 100 µM TGN-020 or 100 µM AER-270 for 10 minutes. Cultured cells were treated with 100 µM TGN-020 or 1 µM AER-270 for 1 h before lysing. Each of the replicates was divided into 16 aliquots, which were subsequently incubated at different temperatures ranging from 45 to 71 °C for 3 minutes. After ultracentrifugation, the soluble protein fractions of each replicate were pooled and analyzed with NanoLC-MS/MS on an Orbitrap Exploris mass spectrometer (Thermo Fisher Scientific). Proteins were identified and quantified against the Uniprot Homo sapiens (Human) protein database UP000005640 or *Xenopus laevis* oocyte database (Taxonomy ID 8355). Only proteins with at least two unique peptides were included in the further analysis. The quantified abundance of each protein in each technical replicate was normalized to the total intensity of the vehicle control to identify proteins with changed thermal stability (solubility) in the presence of the compound. A two-tailed t-test was employed to calculate the p-value. The target candidates for each compound were identified among the proteins with simultaneously large log_10_ p-values in all PISA data.

### Radioligand binding assay

Membrane samples for radioligand binding were prepared to a concentration of 2 mg/mL wet weight of membrane in binding buffer (100 mM sodium chloride, 20 mM HEPES (pH 7.4)). To remove any adenosine from the A_2A_R binding sites, adenosine deaminase was added to a final concentration of 0.1 U/mL.

Various concentrations of unlabelled ZM-241385 were made up in DMSO from 10^-8^ M to 10^-3^ M to yield in assay concentrations of 10^-11^ - 10^-6^ M. TGN-020 was diluted in DMSO to yield a final in assay concentration of 150 µM. The radioligand [^3^H]ZM241385 (American Radiolabelled Chemicals (Royston, UK), specific activity, 80 Ci/mmol) was diluted in binding buffer to yield a final in assay concentration of 80,000 cpm/mL.

For all reactions, 5 µL of [^3^H]ZM-241385 and 500 µL of membrane was added to a tube along with 5 µL of either DMSO, unlabelled ZM-241385 or TGN-020. Membranes were incubated with the ligands for 30 minutes at room temperature then centrifuged at 14,000g for 5 minutes at 4°C. The supernatant was discarded, and the pellets washed twice with water to remove any unbound ligand. The pellets were then air dried for 1 hour. 100 µL of soluene was added to each pellet and incubated at room temperature overnight. Samples were mixed well pre and post addition of 1 mL scintillant before liquid scintillation counting.

### Statistical analysis

Prism version 10 (Graph Pad, San Diego, CA, USA) was used for all analyses. Biological replicates (n) are indicated in the Figure legends and technical replicates are listed in the method section.

## Supplementary figure

**Figure S1.**
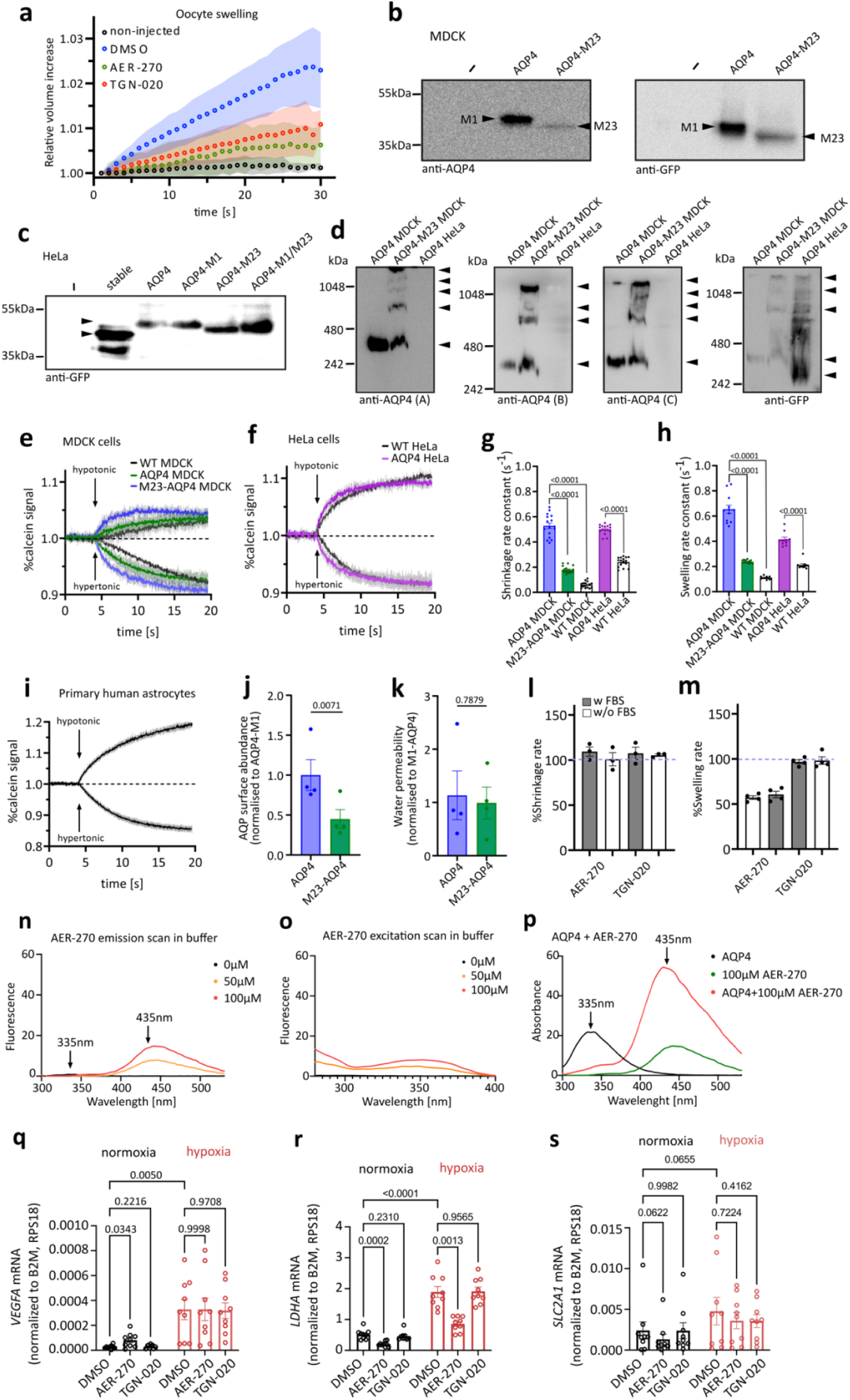
(a) Raw swelling curves of rat AQP4-M1 and control Xenopus laevis oocytes pre-treated with 100 µM AER-270 or TGN-020 (mean ± SD). (b) Western blot analysis of whole cell lysates demonstrated M1 isoform expression in MDCK cells transfected with human AQP4-GFP. and M23 isoform expression in MDCK cells transfected with AQP4-M23-GFP. (c) Stably transfected HeLa cells expressed both human AQP4 isoforms with M23 more abundant. Transiently transfected cells served as a control for molecular weight validation of the two isoforms. (d) Western blot following blue native PAGE demonstrated orthogonal array of particle (OAP) formation in the AQP4-M23 expressing cell lines. (e) Calcein fluorescence quenching demonstrated functional expression of AQP4 in the stable cell lines. Osmotic challenge with hyperosmotic or hypoosmotic solutions led to cell shrinkage (fluorescence signal decrease) and cell swelling (signal increase), respectively, in (e) MDCK and (f) HeLa cells, with higher rates (g, h) in the AQP4-expressing cells compared to the wild-type controls (n>3). (i) Representative calcein fluorescence traces from primary human astrocytes in response to hyperosmotic and hypoosmotic media. (j) ELISA-based cell-surface biotinylation assay for AQP4 membrane abundance in stable MDCK cells (n=4). (k) Shrinkage rates normalised to membrane abundance were not significantly different between the isoforms. (l,m) Water permeability measurements following treatment with 100 µM AER-270 or 100 µM TGN-020 demonstrated no difference in the absence of foetal bovine serum (FBS). (n,o) AER-270 was autofluorescent with an excitation maximum < 280 nm and emission maximum at 435 nm. (p) Increased AER-270 fluorescence signal in the presence of SMA-solubilised AQP4. Hypoxia-responsive control genes (q) LDHA, (r) VEGFA and (s) SLC2A1 (GLUT1) were upregulated in primary human astrocytes exposed to 1% O_2_ for 24 hours, validating the hypoxic setup (n=9). Data are presented as mean ± SEM. Statistical analysis was performed using (g, h) a one-way ANOVA with Bonferroni post hoc test, (j, k) a paired two-tailed t-test or (q-s) two-way ANOVA with Bonferroni post hoc test.

